# Recombinant spider silk protein matrices facilitate multi-analysis of calcium-signaling in neural stem cell-derived AMPA-responsive neurons

**DOI:** 10.1101/579292

**Authors:** Michalina Lewicka, Paola Rebellato, Jakub Lewicki, Per Uhlén, Anna Rising, Ola Hermanson

## Abstract

Neural progenitors or stem cells (NSCs) show great promise in drug discovery and clinical application. Yet few efforts have been made to optimize biocompatible materials for such cells to be expanded and used in clinical conditions. We have previously demonstrated that NSCs are readily cultured on substrates of certain recombinant spider silk protein without addition of animal- or human-derived components. The question remains however whether this material allows differentiation into functional neurons and glia, and whether such differentiation can take place also when the NSCs are cultured within the material in a pseudo-3D context. Here we demonstrate that “foam”-like structures generated from recombinant spider silk protein (4RepCT) provided excellent matrices for the generation and multicellular analysis of functional excitatory neurons from NSCs without addition of animal- or human-derived components. NSCs isolated from the cerebral cortices of rat embryos were cultured on either 4RepCT matrices shaped as foam-like structures without coating, or on conventional polystyrene plates coated with poly-L-ornithine and fibronectin. Upon treatment with recombinant proteins including the growth factor BMP4 or a combination of BMP4 and the signaling factor Wnt3a, the cortical NSCs cultured in 4RepCT foam-like structures differentiated efficiently into neurons that responded to glutamate receptor agonists, such as AMPA, to at least the same extent as control cultures. Matrices derived from recombinant spider silk proteins thus provide a functional microenvironment for neural stem cells without any animal- or human-derived components, and can be employed in the development of new strategies in stem cell research and tissue engineering.

## INTRODUCTION

The microenvironment strongly influences stem cell characteristics in various ways. These effects include changes in differentiation-associated gene expression via epigenetic mechanisms, e.g., chromatin modifications, and related changes in transcriptional activity. Several reports from recent years, including from our lab, have demonstrated how substrate characteristics such as stiffness and roughness, and incorporation of specific cell binding motifs influence the differentiation of various types of stem cells in a cell context-specific manner [1-4]. This is particularly important when addressing neural tissue modeling since three-dimensional context of cell-cell interactions and cytoarchitecture is prohibited in conventional 2D cultures. 3D models of the developing brain, such as cortical organoids, can give insight into interplay between for example neurons and astrocytes, giving rise to functional synapses. Organoid growth relying only on cells self-assembly may however result in mixtures of cells with neural and non-neural characteristics with poor reproducibility, which in turn may lead to non-physiological cell interactions. Accordingly, a wide range of alternative solutions for optimization of stem cell research and tissue engineering has been tested during recent years.

Spider silk has been suggested to be an ideal biomaterial [5-8], due to its extreme mechanical properties [9] combined with local tolerance when implanted in living tissues [10-12]. A recombinant spider silk protein (4RepCT, consisting of 4 tandem repeats and the C-terminal domain of the major ampullate spidroin 1) that corresponds to approximately 10% of the native spider silk protein, can be readily produced in *E.coli* and spontaneously assembles into films, foams and fibers [13, 14]. 4RepCT matrices provide excellent substrates for human primary fibroblasts [14], and we have demonstrated that this recombinant spider silk protein provides a suitable substrate also for culturing of rodent cortical neural stem cells (NSCs) as these cells proliferate, survive, retain multipotency, and differentiate efficiently on film structures derived from 4RepCT [15]. The analysis of differentiation in this previous study was based on gene expression and protein production, and it is still not known whether the differentiated cells derived from neural stem cells grown on 4RepCT are functional. As it has been shown repeatedly that the expression of differentiation markers not necessarily correlate with functionality [16], such functional analysis is required before proceeding with transplantations and *in vivo* studies in animals.

## MATERIALS AND METHODS

### Production of protein matrices

The recombinant miniature spider silk protein 4RepCT was produced in *Escherichia coli* and purified as described previously [17]. A procedure for depletion of lipopolysaccharides (LPS) was also included [18]. After purification, the protein solution was sterilized by passage through 0.22 μm filter and concentrated to 3 mg/mL by ultrafiltration (Amicon Ultra, Millipore) before preparation of foam matrices according to Widhe *et al* [14]. Matrices were allowed to dry over night at room temperature under sterile conditions and stored in room temperature until use. Before plating the cells, the matrices were washed twice with sterile PBS and pre-incubated with serum-free DMEM:F12 media (Sigma-Aldrich) for 1 h at 37 °C with 5% CO_2_. The matrices were prepared in six-well cell culture polystyrene plates or 35mm in diameter polystyrene plates (Sarstedt). For surface morphology, SEM and in depth analysis of the substrates, please see Widhe et al [14].

### Neural Stem Cell (NSC) culture

*N*SCs were isolated from the cerebral cortices of day E15.5 embryos obtained from pregnant Sprague Dawley rats largely as described previously [16, 19]. Cells were mechanically dissociated followed by culture in serum-free DMEM:F12 media (Gibco) enriched with N2 supplements. Cells were maintained in proliferating state using 10 ng/ml fibroblast growth factor 2 (FGF2; R&D Systems) and passaged twice before usage in the experiments. NSCs were then seeded at a density of 10,000 cells/cm^2^ and allowed to proliferate for 24-72 hrs prior to the experiments. The 4RepCT foams covered ~60% of the surface area of the cell culture dishes before NSCs were applied. NSCs used in positive control experiments were seeded on conventional polystyrene culture plates coated with poly-L-ornithine and fibronectin from bovine plasma (P+F) (Sigma-Aldrich) to which uncoated polystyrene dishes served as negative controls. Soluble factors were re-supplemented at 24 hrs and medium replaced at 48 hrs time intervals. To induce differentiation FGF2 was withdrawn from the cell culture media and then replaced with bone morphogenic protein (BMP4; R&D Systems) or and Wnt3a (R&D Systems). All animal experiments were approved by the North Stockholm regional committee for animal research and ethics, Stockholm, Sweden (N284/11).

### Immunocytochemistry

Cell cultures were first rinsed once in Phosphate-Buffered Saline (PBS) (Gibco) and then fixed in 10% Formalin (Sigma-Aldrich) for 20 min at room temperature (RT) followed by 3 x 5 min washes with PBS/0,1% Triton-X 100 (Sigma-Aldrich). Plates were then incubated with the respective primary antibody in PBS/0,1% Triton-X 100/1% bovine serum albumin (BSA; Sigma-Aldrich) overnight at 4°C. The primary antibodies sources and dilutions were as follows: rabbit poly-clonal anti-glia fibrillary acidic protein (GFAP; 1:500; DAKO), mouse monoclonal anti-Neuronal Class III B-Tubulin (TuJ1; 1:500; Covance), mouse anti-intermediate filament protein nestin (1:500; BD Pharmingen), microtubule associated protein 2 (MAP2) (1:500, Sigma-Aldrich). The cells were subsequently washed 6 x 5 min each with PBS/0,1% Triton-X 100. Secondary antibodies were incubated in PBS/0,1% Triton-X 100/ 1% BSA at room temperature for 1 hr. The secondary antibodies were species-specific labeled with Alexa-488 or Alexa 594 (1:500; Molecular Probes). Next the samples were washed 3 x 5 min each in PBS and mounted with Vectashield (Vector Laboratories, Inc). Fluorescent images were acquired with Axioskop software using Zeiss Axioskop2 microscope coupled to an MRm (Zeiss).

### Calcium imaging

Cells were loaded with the Ca^2+^sensitive fluorescence indicator Fluo-3/AM (5 µM, Life Technologies) together with 0.1% Pluronic f-127 (Life Technologies) at 37°C for 20 min in their own medium, and then moved to a Krebs-Ringer buffer containing (in mM) 119.0 NaCl, 2.5 KCl, 2.5 CaCl_2_, 1.3 MgCl_2_, 1.0 NaH_2_PO_4_, 20.0 HEPES (pH 7.4), and 11.0 dextrose. Measurements of intracellular Ca^2+^ were carried out in Krebs-Ringer buffer with or without Ca^2+^ at 37°C using a heat-controlled chamber (Warner Instruments) with a cooled EMCCD QuantEM 512:SC camera (Photometrics) mounted on an upright microscope (Zeiss Axio Examiner.D1) equipped with an W-Plan apochromat 20X/1.0 objective (Carl Zeiss). Excitation at 495 nm was assessed with a filter wheel (Sutter Instrument) at sampling frequency 0.5 Hz. MetaFluor (Molecular Devices) was used to control all devices and to analyze acquired images. The response to AMPA and ATP was determined using MATLAB software (The MathWorks Inc.) as described previously [20] considering only increases of more than 30% compared with the baseline as positive response. Drugs were bath-applied following previous protocol [16].

### Statistical analysis

Statistical analysis and graphs were performed using the software Prism 4 (Graph Pad). When 2 groups were compared, statistical analysis was performed using two-tailed unpaired t-test. When repeated measurements taken at different time points from 2 groups were compared, two-way ANOVA analysis of variance was used. The threshold value for statistical significance (*α* value) was set at 0,05 (*p < 0,05). Data are presented as the mean ± standard error of the mean (SEM) of independent experiments.

## RESULTS

### Control cultures

In control cultures, NSCs derived from rat cortex of embryonic day 15 (E15) were cultured in optimal conditions seeded on polystyrene cell culture plates precoated with poly-L-ornithine and fibronectin. This protocol is well-established and widely used, and provides homogenous cultures of multipotent progenitors for a large number of passages [2, 21-24]. The NSCs expand undifferentiated and self-renew in N2 supplemented medium in the presence of FGF2. In contrast, NSCs grow poorly or not at all on uncoated polystyrene plates or plates coated with either poly-ornithine or fibronectin [2, 15, 21].

### 4RepCT foam structures provide efficient substrates for neuronal differentiation of neural stem cells as assessed by morphology and markers

After initial expansion of the primary cells, we seeded NSCs after the first passage on 4RepCT foam structures and in control conditions, respectively. NSCs seeded on 4RepCT foam structures attached and proliferated 24-72 hrs after the passage (**Figure 1**), indicating that NSC can attach to the 4RepCT proteins as previously reported. To investigate if NSCs grown on 4RepCT foam shaped matrices in 3D can differentiate into neurons we exposed the cells to treatment with bone morphogenetic protein (BMP) 4 or co-treatment with BMP4 and the signaling factor Wnt3a for 14 days. We have previously shown that BMP4 and BMP4+Wnt3a stimulation induces differentiation of mature, functional glutamate-receptor (GluR) –agonist responsive neuronal cells when NSCs are seeded at relatively high densities [16, 25]. NSCs grown in 3D on 4RepCT foam matrices with pore sizes of around 5-30 µm and exposed to 10 ng/ml BMP4 or BMP4 and Wnt3a every 24 hours for 14 days differentiated into neuronal cells positive for the dendritic marker of mature, post-mitotic neurons, microtubule associated protein 2 (MAP2; in average 30 cells/5 micrographs/well (BMP4) and 41 cells/5 micrographs/well (BMP4+Wnt3a) in control conditions, and 71 cells/5 micrographs/well (BMP4) and 94 cells/5 micrographs/well (BMP4+Wnt3a) when grown on 4RepCT foam) (**Figure 1,2**). Morphological characteristics of cells grown on 4RepCT foam structures were similar to the poly-ornithine and fibronectin-coated polystyrene cell culture plates used as controls (**Figure 1**). Virtually all MAP2-negative cells were found to stain positive for the astrocytic marker GFAP (glial fibrillary acidic protein) (**Figure 1**). NSCs grown on 4RepCT 3D foam-like structures thus retained full potential to differentiate into cells expressing neuronal and astrocytic markers, and 4RepCT-derived structures seemed to provide an even better substrate for differentiation than standard culture plates as assessed by MAP2 staining.

**Figure 1.**
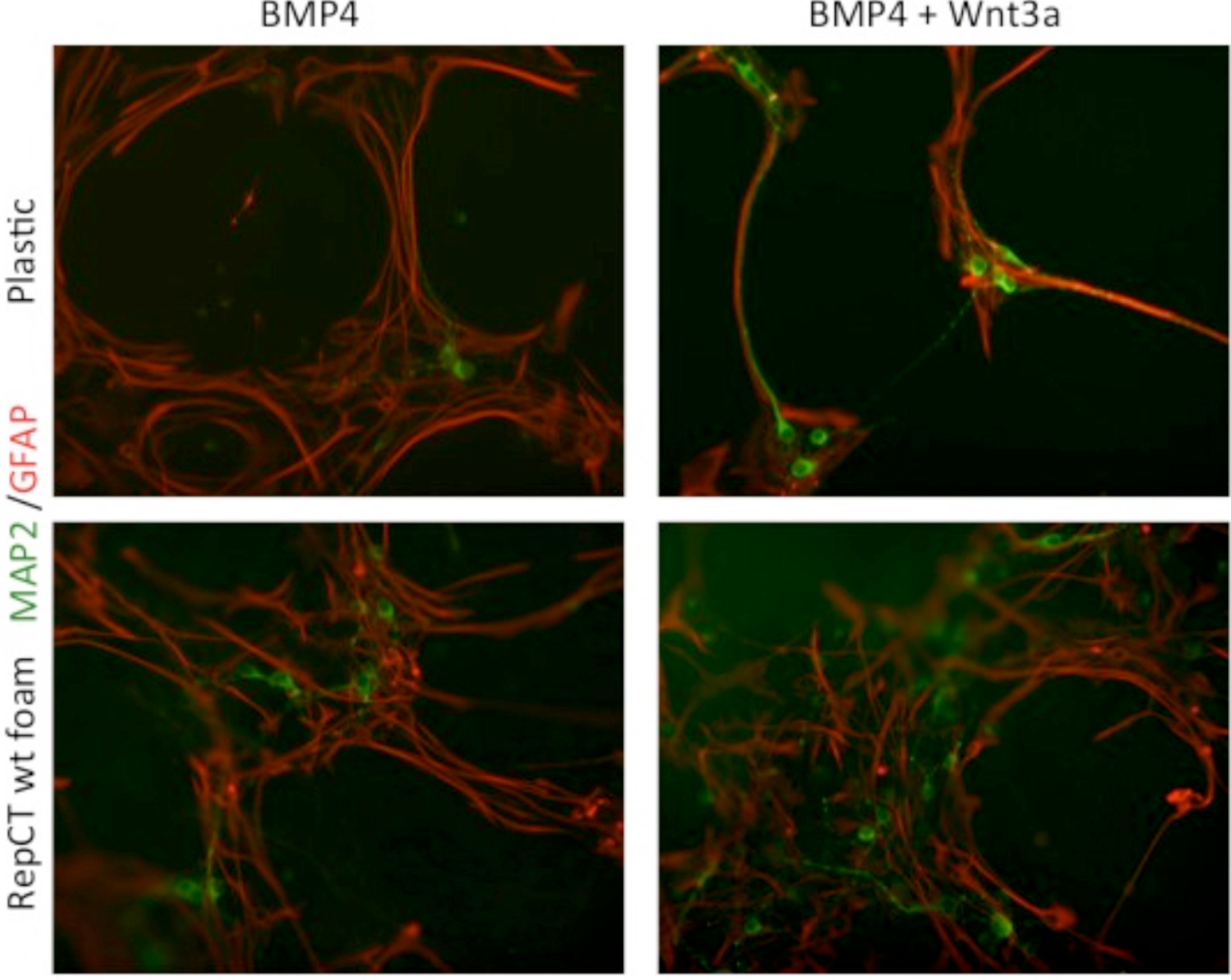
NSCs retained their potential to differentiate into mature post-mitotic neuronal cells on 4RepCT films in response to BMP4 and co-treatment of BMP4 and Wnt3a respectively. Micrographs demonstrating neuronal cells derived from NSCs after treatment with BMP4 only (A,B) or co-treatment with BMP4 and Wnt3a (C,D) when grown on control culture plates (A,C) or 4RepCT (B,D). Green=MAP2-staining. Red=GFAP-immunoreactivity. Blue=DAPI.

**Figure 2.**
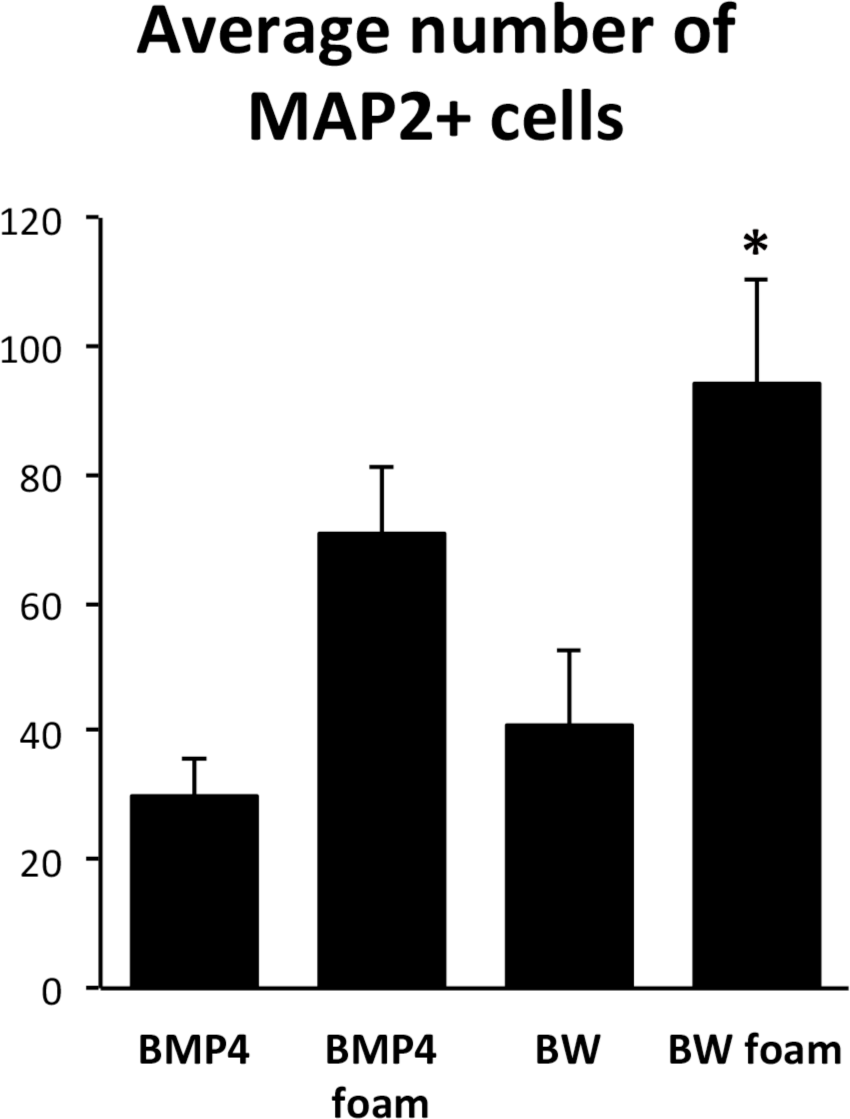
Quantifications of the fractions of MAP2-stained cells relative to the number of DAPI stained cells after BMP4 or co-treatment with BMP4 and Wnt3a (BW) on the 4RepCT-derived foam matrix. Y axis represents average number of MAP2-positive cells in 5 micrographs similar to those shown in Figure 1 from three independent experiments in each condition. *=p<0.05, error bars represent SEM.

### 4RepCT foam structures provide efficient substrates for differentiation of neural stem cells into AMPA-responsive, functional neurons

BMP4 and Wnt3a are required for proper development of neurons of the hippocampal formation, a structure of the forebrain essential for learning, and mature hippocampal neurons express receptors for the excitatory neurotransmitter glutamate [26, 27]. We therefore decided to investigate the functionality of BMP4 versus BMP4+Wnt3a induced neurons grown in 3D 4RepCT matrices by examining their responsiveness to specific glutamate receptor and calcium (Ca^2+^) signaling agonists as previously described [16]. Calcium signaling was measured in NSCs differentiated for 14 days and loaded with the Ca^2+^ sensitive fluorophore Fluo-3/AM.

All cells examined in BMP4 or BMP4+Wnt3a-treated cultures were first shown to respond to KCl depolarization with a rapid increase in the intracellular Ca^2+^ concentration (data not shown), indicating that the cultures were functional and healthy. To investigate the functional identity of the cells, we used an established protocol to discriminate between neurons and astrocytes in the BMP4- and BMP4+Wnt3a-treated cells. Astrocytes express mainly the metabotropic P2Y purinoceptors whereas mature neurons express the ionotropic P2X purinoceptors, plus a small population of P2Y receptors that do not affect intracellular Ca^2+^ but mediate slow changes in membrane potential [28, 29]. Neurons should therefore respond to ATP in the presence, but not absence, of extracellular Ca^2+^, whereas astrocytes should respond to ATP in both conditions [28, 30]. Cells were further characterized functionally by treatment with the selective ionotropic glutamate receptor (GluR) agonist AMPA, which, in the presence of extracellular Ca^2+^, should evoke a transient response in neurons but not in astrocytes. Stimulation of BMP4 and BMP4+Wnt3a-treated cultures with ATP or AMPA in an extracellular medium with or without Ca^2+^ confirmed our previous observations that the NSC-derived cells can be divided into several subsets depending on their distinct Ca^2+^ response patterns. One subset of cells responded to ATP and AMPA, respectively, with a transient Ca^2+^ increase in the presence but not absence of extracellular Ca^2+^, while another subset of cells only responded to ATP but not AMPA under both conditions. Typical response curves are depicted in **Figure 3**. **Supporting Information** contains 4 micrograph time-lapse movies of around 90 seconds each representing one c:a 60-70 minute experiment in each condition demonstrating the calcium response to ATP, AMPA, and KCl similar to the results from each condition shown in **Figure 3.**

**Figure 3.**
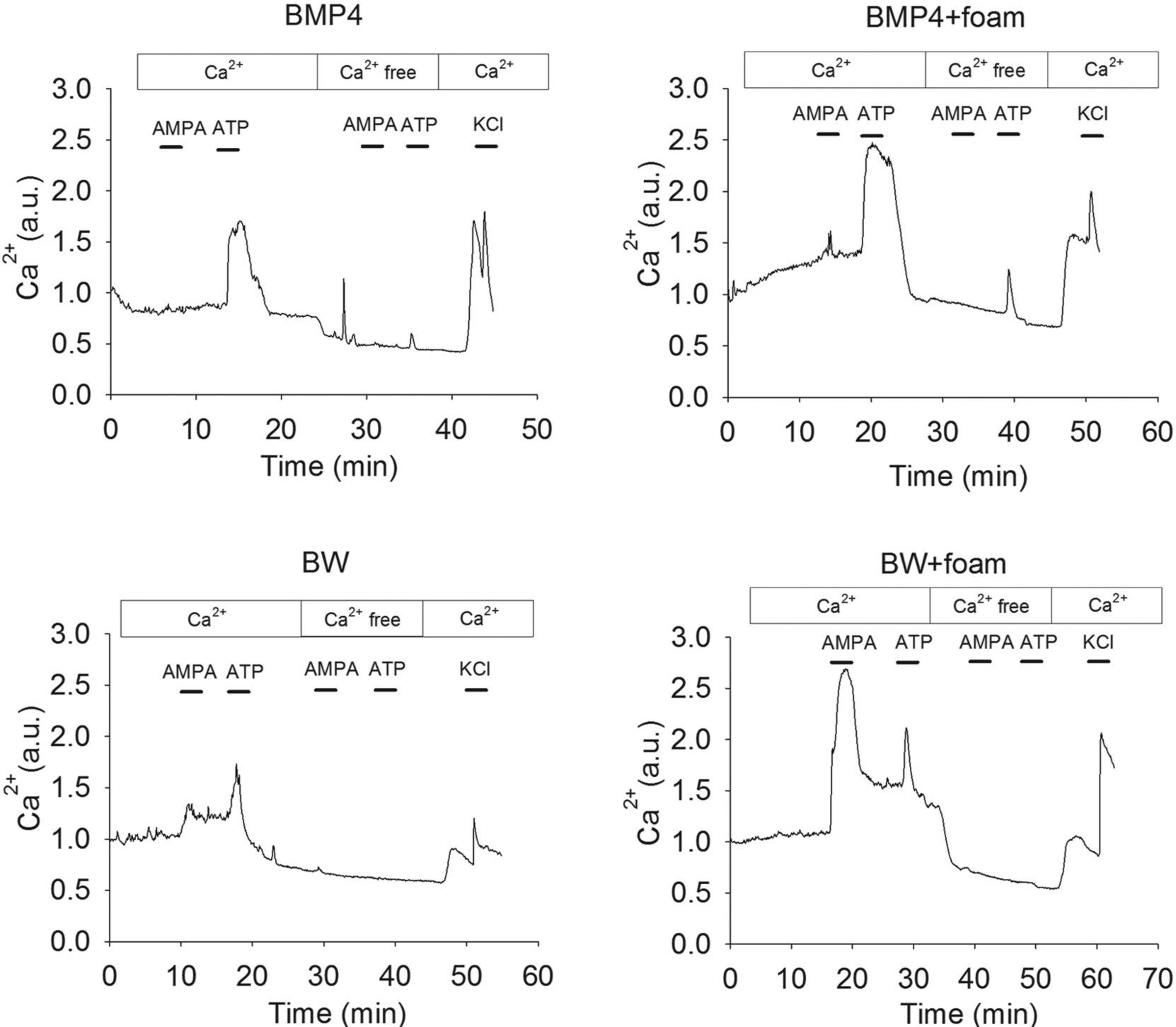
Representative single cell Ca^2+^ response traces after treatment of NSC cultured in control or 4RepCT-derived foam structures with BMP4 and BMP4+Wnt3a (BW).

### 4RepCT foam structures enhance the differentiation of neural stem cells into AMPA-responsive, functional neurons

We next performed a quantitative analysis on BMP4- and BMP4+Wnt3A-treated NSC cultures and their ability to respond to AMPA. This analysis revealed that AMPA stimulation resulted in a transient Ca^2+^ increase in ≈25% of the cells in NSC-derived cultures grown in 3D 4RepCT foams and differentiated by co-treatment of BMP4 and Wnt3a compared to around 10% grown on control substrates (**Figure 4**). The functional characterization of NSC-derived neurons validated our previous observation [16] that expression of markers for post-mitotic neurons, e.g., MAP2, were not sufficient to make conclusions regarding the functionality of stem cell-derived neurons.

**Figure 4.**
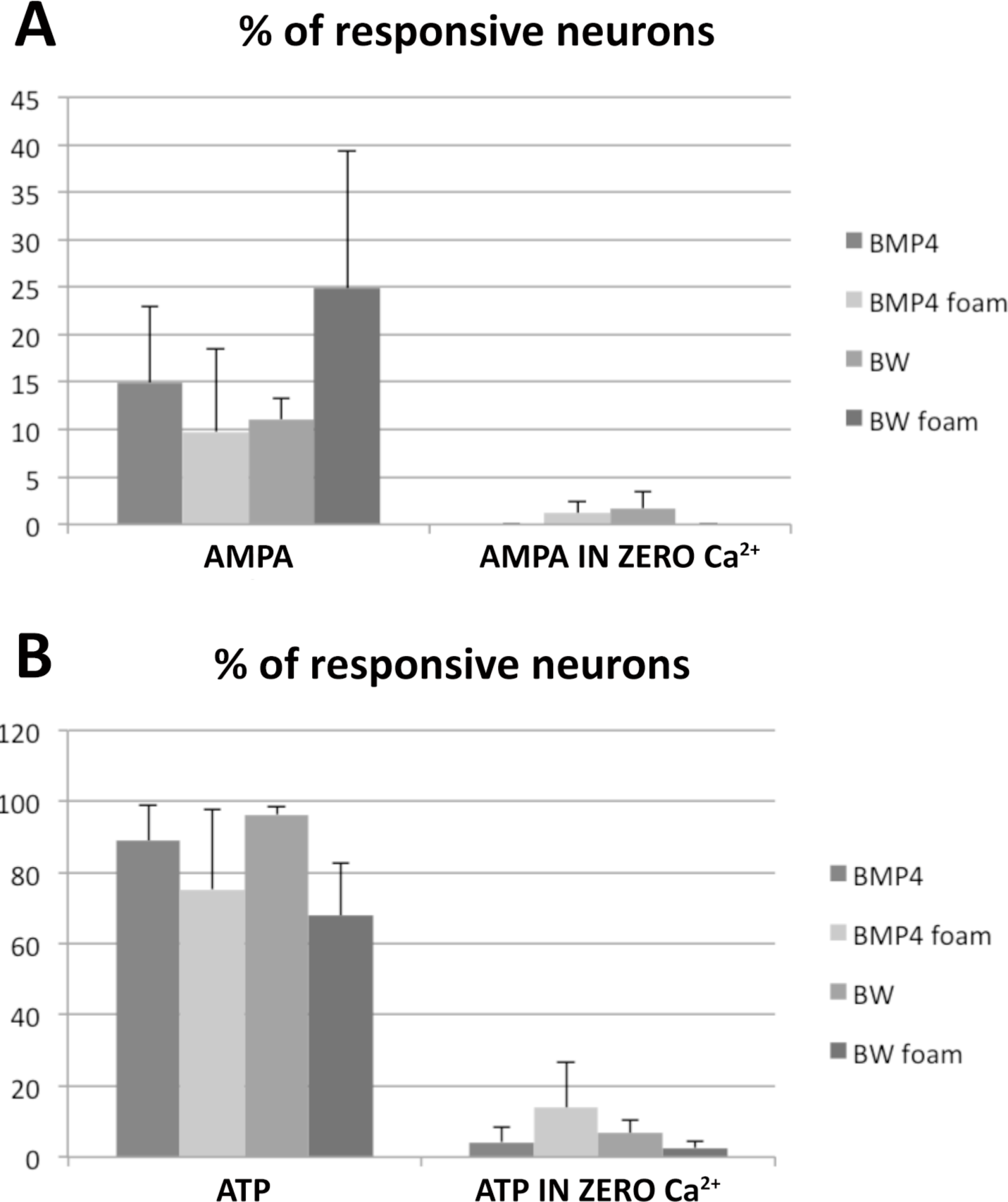
Quantifications of the Ca^2+^ imaging experiments. A large variability was noted, but the largest fraction of AMPA-responsive neurons was clearly found in the BMP4+Wnt3a (BW)-treated cultures grown in 3D 4RepCT-matrices. Each bar represents the means ± SEM (n>30 cells per condition) of single cell Ca^2+^ responses as illustrated in Figure 3 from n=3 independent experiments.

## DISCUSSION

So far, most approaches in cell therapy of the central nervous system (CNS) have been based on administering relatively homogenous cells to rather homogenous brain structures. However, CNS is highly organized with specific functional centers consisting of mixed cell types, and due to the strictly organized anatomy of the CNS, 3D-approaches are not only recommended, it is required for continued progress in applied stem cell research and regenerative medicine. Recombinant spider silk provide several potential advantages over other materials used as matrices and/or over the natural material: (i) it displays mechanical integrity and can be processed into different 3D structures [13, 14], (ii) the production is scalable [31], (iii) the sequence of the proteins can easily be modified to contain also cell binding motifs [32], (iv) recombinant spider silk fibers evoke little inflammatory reaction and are gradually degraded when implanted subcutaneously [33], and (v) the material is chemically defined and of non-animal origin, and thus a possible substrate for long-term, xeno-free cell culture.

Our system has some differences and advantages compared to most published protocols hitherto. First, as previously described, NSCs cultured and differentiated on 4RepCT do not require any additional coating, allowing a xeno-free substrate for organoid cultures as mentioned. In addition, the 4RepCT substrate is transparent allowing analysis of morphological details. Due to this transparency, we could perform calcium analysis after agonist stimulation, and demonstrate the overall efficiency of the differentiation and nature of the differentiated cells and structures of a large number of cells. Furthermore, in contrast to many but not all other protocols, our rodent cells have been defined as multipotent and able to differentiate into neurons, astrocytes, and oligodendrocytes in clonal conditions [23] as well as mesenchymal fates allowing a more strict control of the development of the tissue-like structures. Lastly, as the spider silk substrate consists of recombinant protein, it opens up for various possibilities of developing simple protocols for growing neural organoids in GMP-grade environment in the near future.

Here we employed 4RepCT foams in NSCs differentiation in order to successfully identify a defined 3D culture system for NSCs. The 4RepCT foam structure supports NSC attachment, survival, and differentiation into functional neurons in 3D, and 4RepCT 3D foam implantation has already been successfully applied in animal studies, suggesting it to be a near-ideal candidate for future studies of NSC differentiation and formation into functional circuits *in vitro* and for tests for use of NSCs in cell therapy in various models of neurological disease *in vivo*. Current studies focus on the implementation of multipotent human induced pluripotent stem cell (hIPSC)-derived neural progenitors and 3D bioprinting to achieve optimally controlled development of human brain/ganglion-like structures *in vitro*.

## CONCLUSION

Foam-like structures generated from recombinant spider silk protein (4RepCT) permit differentiation of neural stem cells into functional, AMPA-responsive neuronal circuits in 3D without animal- or human-derived components. We propose that matrices generated from recombinant spider silk protein are suitable for development of applications in stem cell research, tissue engineering, and regenerative medicine.

## Abbreviations

NSCs: (neural stem cells)
BMP4: (Bone Morphogenic protein 4)
TuJ1: (Neuronal class III B-tubulin)
P: (poly-ornithine)
F: (fibronectin)
ECM: (extracellular matrix)
3D: (three dimensional)
BW: (BMP4+Wnt3a)

## ACKNOWLEDGEMENTS

The authors would like to thank the Hermanson lab for valuable discussions and comments on this manuscript. We thank Spiber Technologies AB for providing the 4RepCT matrices. This work was funded by grants from Karolinska Institutets Forskningsstiftelser (2011FoBi023) (to A.R.), VR-MH (project grants to P.U. and O.H.), Karolinska Institutet (TEMA), the Swedish Childhood Cancer Foundation (BCF), the Swedish Cancer Society, and Vinnova (to O.H.).

